## Supplementary Figures for "Phytochrome-dependent responsiveness to root-derived cytokinins enables coordinated elongation responses to combined light and nitrate cues"

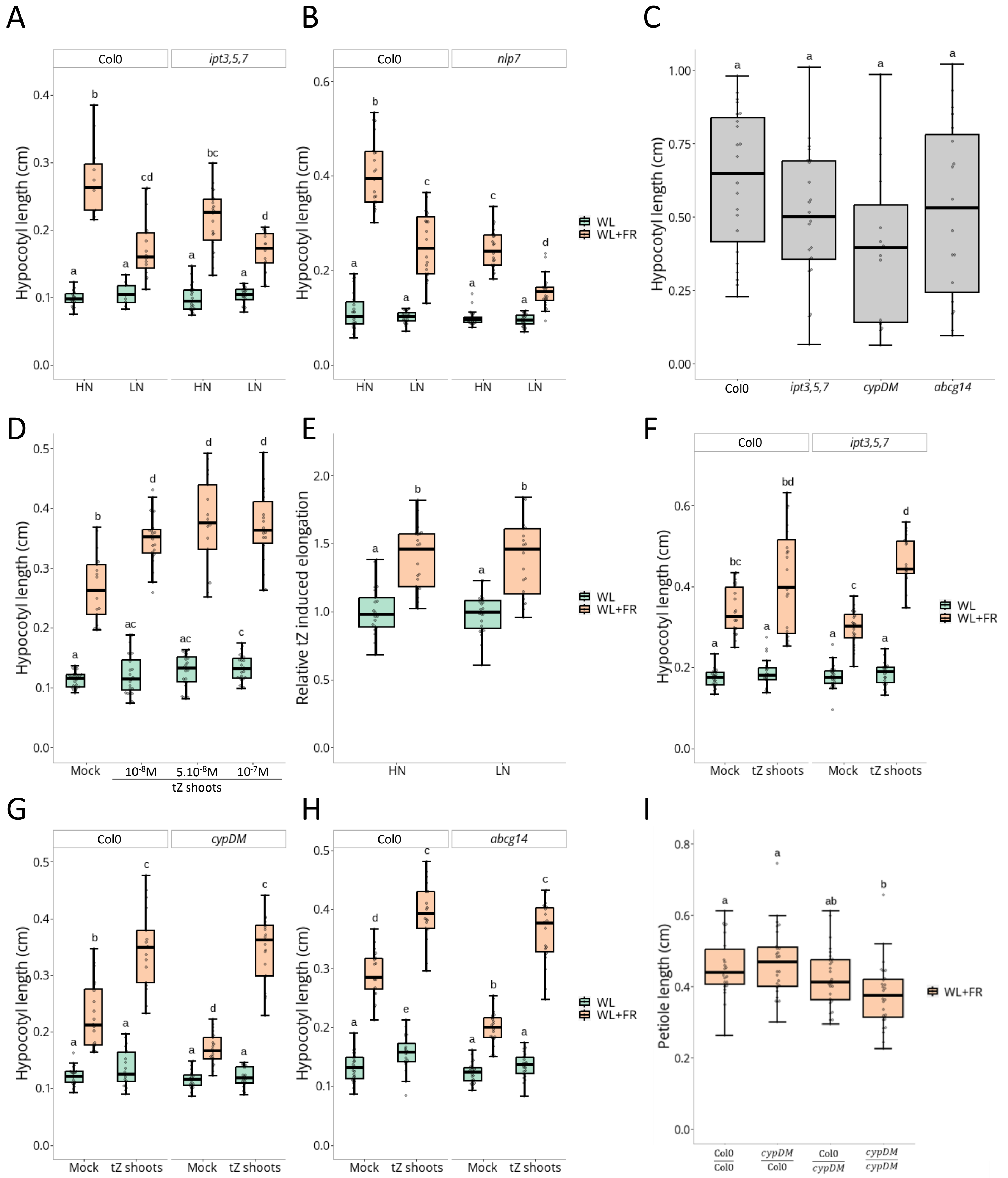


Sup. Figure 1: transZeatin modulates FR light-induced hypocotyl elongation in response to nitrate

Sup. Figure 1: transZeatin modulates FR light-induced hypocotyl elongation in response to nitrate

(**A**) Hypocotyl length in cm of Col-0 and *ipt3,5,7* (n>10 plants per condition) or *nlp7* (**B**, n>22) seedlings grown on High Nitrate (HN, 10 mM) or Low Nitrate (LN, 0.2 mM) for 4 days under White Light (WL) and then transferred 4 more days to WL or White Light + Far-Red light (WL+FR). (**C**) Hypocotyl length in cm of Col-0, *abcg14*, *ipt3,5,7* and *cypDM* mutants germinated in the dark for 3 days (n>15). (**D**) Hypocotyl length in cm of Col-0 seedlings grown for 4 days under WL and then transferred 4 more days to compartment plates treated with Mock, tZ $10^{-8}$ M, $5.10^{-8}$ M, or $10^{-7}$ M on the shoot compartment and under WL or WL+FR (n>16). (**E**) Associated to Figure 1G. Relative tZ induced elongation for each tZ treated plant, compared to the mean of all Mock treated plants per condition. (**F**) Hypocotyl length in cm of Col-0 and *ipt3,5,7* (n>21); *cypDM* (**G**, n>16) or *abcg14* (**H**, n>19) seedlings grown for 4 days under WL and then transferred 4 more days to compartment plates treated with Mock or tZ ($10^{-8}$ M) on the shoot compartment and under WL or WL+FR. (**I**) Petiole length in cm of hypocotyl-grafted Col-0 and *cypDM* plants after a ~10days recovery period and treated 4 days with WL+FR (n>18). Different letters depict statistical differences according to a Kruskal-Wallis test (p<0.05).

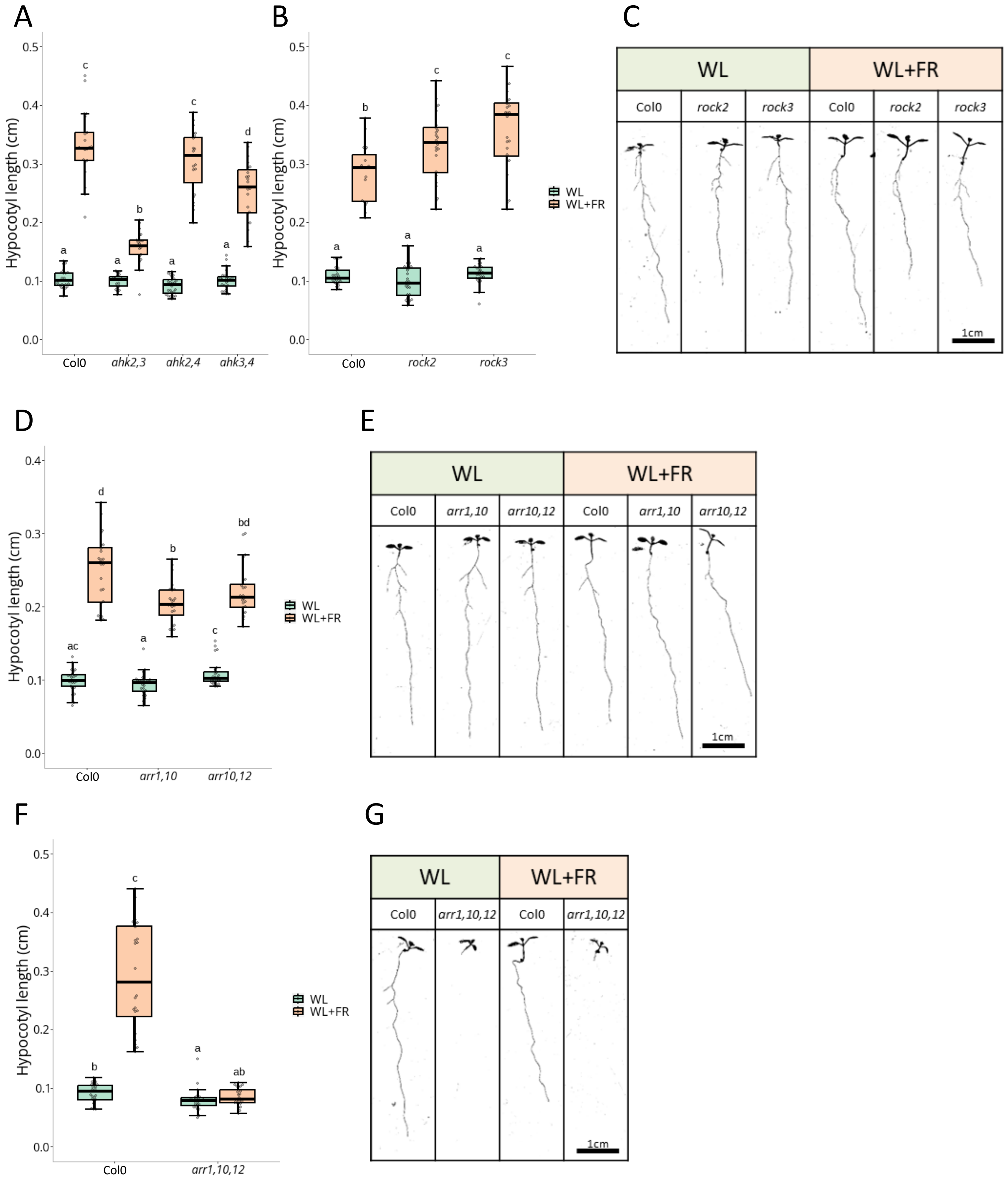


Sup. Figure 2: Cytokinin signalling actors are involved in shade avoidance responses

Sup. Figure 2: Cytokinin signalling actors are involved in shade avoidance responses

(**A**) Hypocotyl length in cm of Col-0 and *ahk2,3, ahk2,4, ahk3,4* (n>17 plants per condition); *rock2, rock3* (**B**, n>18, representative images in **C**); *arr1,10, arr10,12* (**D**, n>23, representative images in **E**) or *arr1,10,12* (**F**, n>22, representative images in **G**) grown for 4 days under White Light (WL) and then transferred 4 more days to WL or White Light + Far-Red light (WL+FR). Different letters depict statistical differences according to a Kruskal-Wallis test ($p<0.05$).

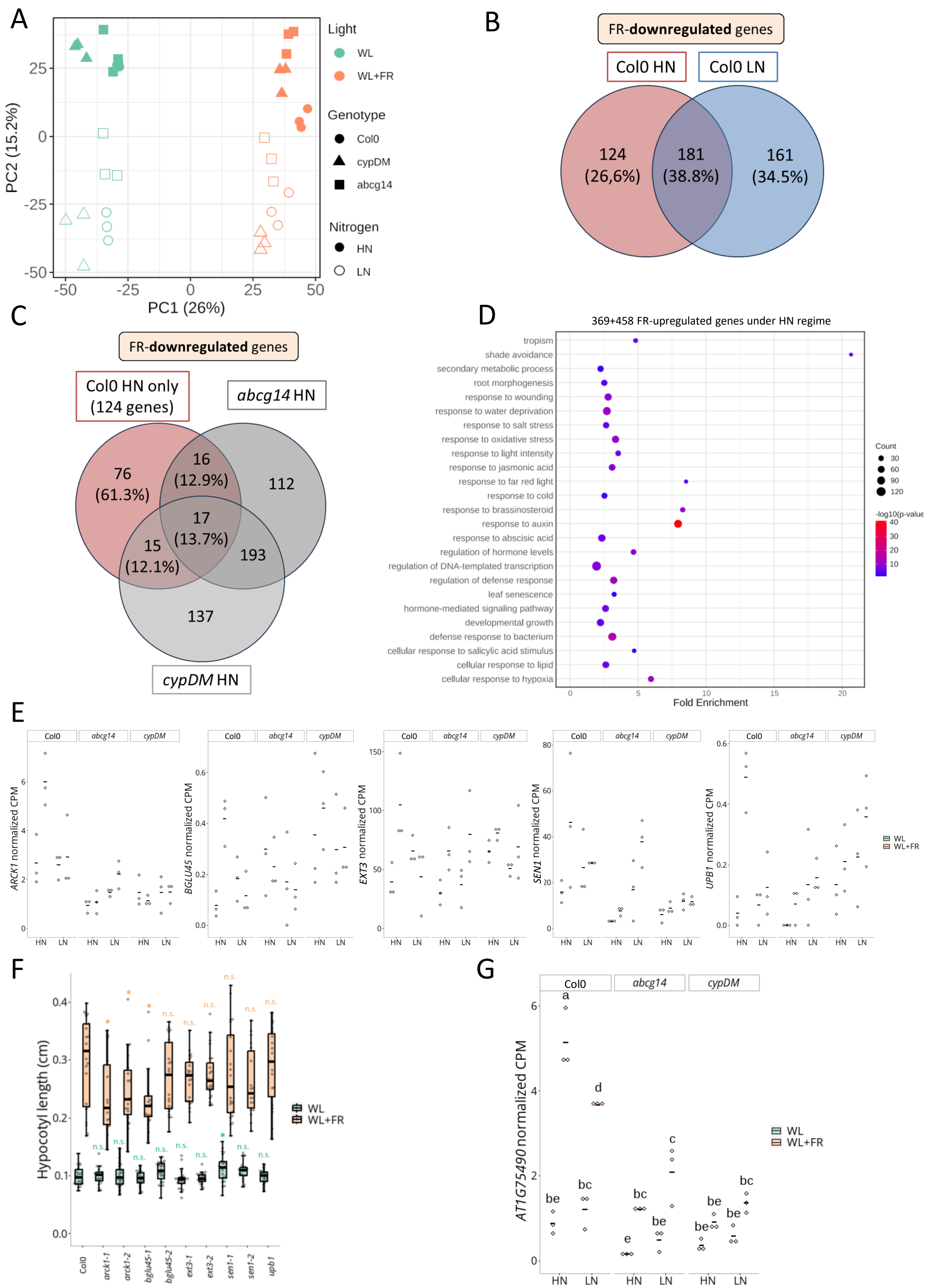


Sup. Figure 3: Nitrate availability and tZ deficiency widely affects FR light-induced transcriptome changes, leading to the identification of a novel shade avoidance actor

Sup. Figure 3: Nitrate availability and tZ deficiency widely affects FR light-induced transcriptome changes, leading to the identification of a novel shade avoidance actor

(**A**) Principal Component Analysis (PCA) plot. X axis represents PC1 and Y axis PC2. Light conditions are highlighted by different colours, White Light (WL) and White Light + Far-Red light (WL+FR); genotypes with different shapes, Col-0, *acbg14* and *cypDM*; and nitrate conditions with plain or empty shapes, High Nitrate (HN, 10 mM) or Low Nitrate (LN, 0.2 mM). (**B**) Venn diagram representing the overlap of Col-0 HN vs Col-0 LN genes downregulated by WL+FR compared to their respective WL controls ($log_2$ FC≥1, FDR<0.05). (**C**) Venn diagram representing the overlap of the 124 genes only downregulated by WL+FR in Col-0 under HN conditions and genes downregulated in *cypDM* and *abcg14* by WL+FR compared to their respective WL controls ($log_2$ FC≥1, FDR<0.05). (**D**) Bubble plot representing GO enrichment analysis for the 458+369 genes upregulated by WL+FR under HN conditions in Col-0. For GO BP, the most specific category subclasses with a significant enrichment (Fisher's Exact test, Bonferroni corrected, p<0.05) are plotted. X axis represents the fold enrichment compared to a random sample of genes. -$log_{10}$(p-value) is indicated by colours and number of genes per category is indicated by the dots size. (**E**) Normalized Count Per Million (CPM) values of *ARCK1, BLUG45, EXT3, UPB1* and *SEN1* across all transcriptome samples. Each dot represents a biological replicate (pool of n>20 plants) and black bars the mean of the biological replicates. No condition combination is upregulated by WL+FR compared to their respective WL controls, except Col-0 HN ($log_2$ FC≥1, FDR<0.05). (**F**) Hypocotyl length in cm of Col-0, *arck1-1*, *arck1-2*, *bglu45-1, bglu45-2, ext3-1*, *ext3-2*, *sen1-1, sen1-2, and upb1* (n>16) grown for 4 days under White Light (WL) and then transferred 4 more days to WL or White Light + Far-Red light (WL+FR). Asterisks depict statistical differences according to a Mann-Whitney test (p<0.05), compared to the respective Col-0 WL (green) and Col-0 WL+FR (orange) controls. (**G**) Normalized Count Per Million (CPM) values of *AT1G75490* across all transcriptome samples. Each dot represents a biological replicate (pool of n>20 plants) and black bars the mean of the biological replicates. Different letters depict significant differences according to a two-way ANOVA followed by a Tukey's post hoc test (p<0.05).

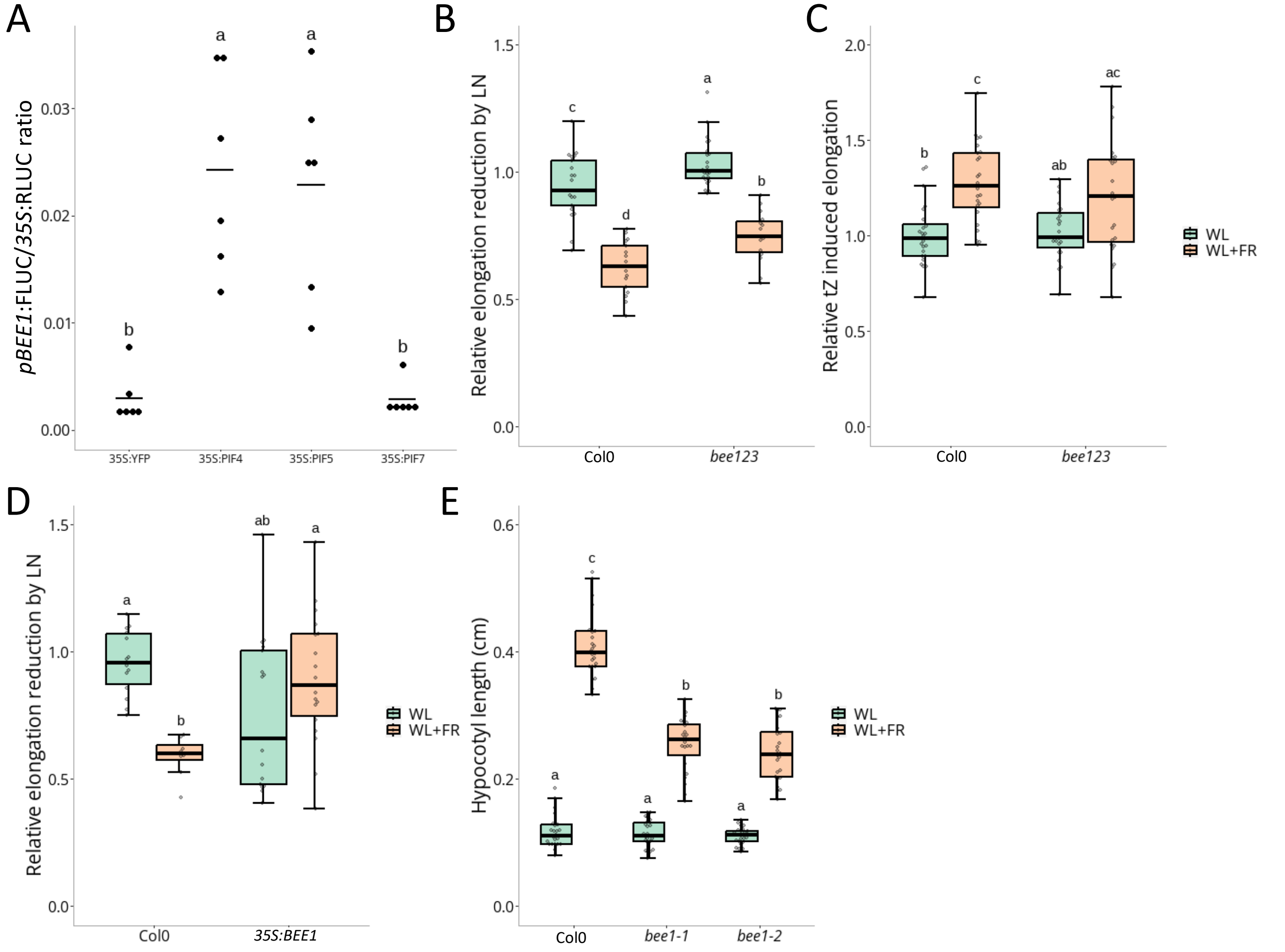


Sup. Figure 4: PIF4 and PIF5 activate *BEE1* expression, and *BEE1* is an integrator of shade and nitrate cues

<u>Sup. Figure 4: PIF4 and PIF5 activate *BEE1* expression, and *BEE1* is an integrator of shade and nitrate cues</u>

(**A**) Transactivation assay in *Nicotiana benthamiana*. A construct expressing *pBEE1:FireflyLUC* and *p35S:RenillaLUC* was co-infiltrated with a construct expressing *p35S:YFP* (baseline control) or *p35S:PIF4, p35S:PIF5* or *p35S:PIF7*. The FireflyLUC reporter activity was expressed ratiometrically to the RenillaLUC internal control. Each dot represents a biological replicate (n=6) and black bars the mean of the biological replicates. The letters depict significant differences according to a two-way ANOVA followed by a Tukey's post hoc test ($p < 0.05$). (**B**) Associated to Figure 3B. Relative elongation reduction by LN for each LN grown plant compared to the mean of all HN grown plants per condition. (**C**) Associated to Figure 3C. Relative tZ induced elongation for each tZ treated plant, compared to the mean of all Mock treated plants per condition. (**D**) Associated to Figure3D. Relative elongation reduction by LN for each LN grown plant compared to the mean of all HN grown plants per condition. (**E**) Hypocotyl length in cm of Col-0, *bee1-1* and *bee1-2* seedlings grown for 4 days under White Light (WL) and then transferred 4 more days to WL or White Light + Far-Red light (WL+FR, n>22 plants per condition). Different letters depict statistical differences according to a Kruskal-Wallis test ($p<0.05$).

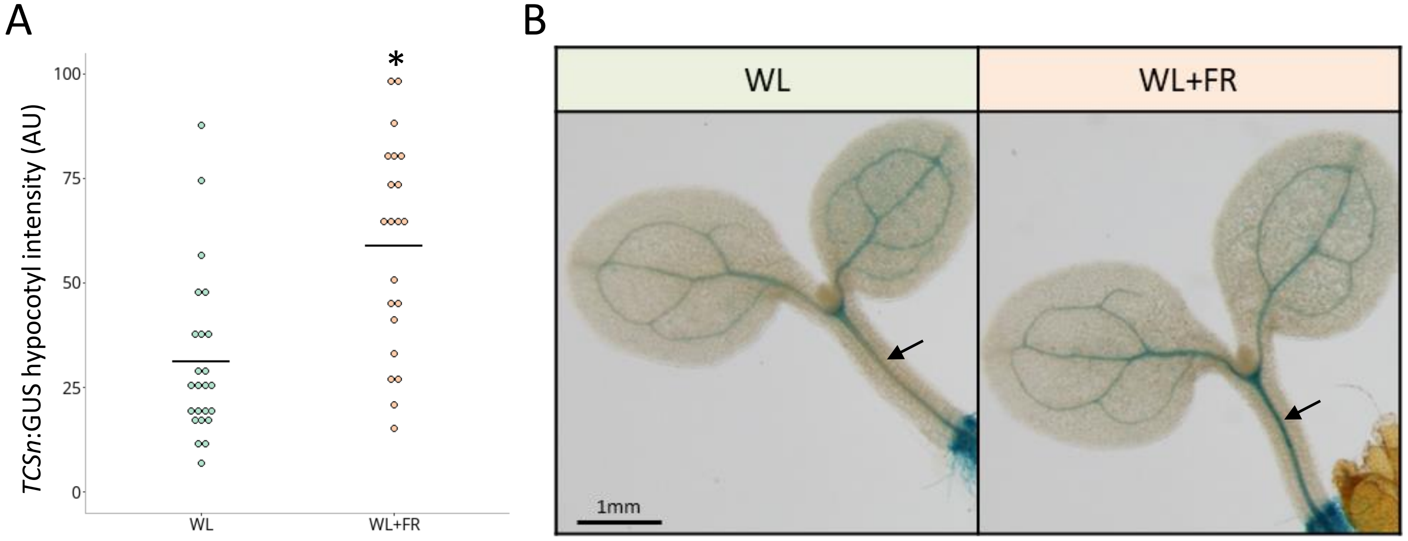


Sup. Figure 5: CK signalling increases in the hypocotyl in response to WL+FR

Sup. Figure 5: CK signalling increases in the hypocotyl in response to WL+FR

**(A)** GUS intensity (AU) in the hypocotyl of *TCSn*:GUS seedlings grown for 4 days under White Light (WL) and then transferred for 90 minutes to WL or White Light + Far-Red light (WL+FR). Each dot represents a biological replicate (n>20) and black bars the mean of the biological replicates. The asterisk depicts a significant difference according to a two-way ANOVA followed by a Tukey's post hoc test ($p<0.05$). (**B**) Representative images of shoots from seedlings in (**A**). Black arrows highlight the GUS signal in the hypocotyl vasculature.

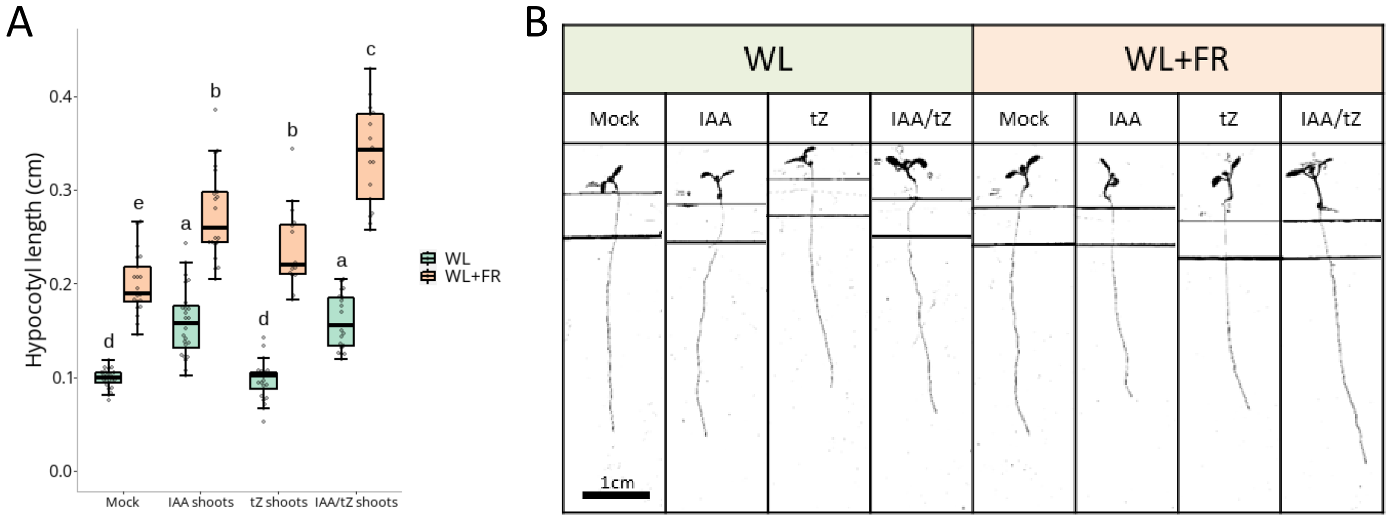


Sup. Figure 6: Auxin alone is not sufficient to potentiate tZ positive role on hypocotyl elongation

<u>Sup. Figure 6: Auxin alone is not sufficient to potentiate tZ positive role on hypocotyl elongation</u>

(**A**) Hypocotyl length in cm and representative images (**B**) of Col-0 seedlings grown for 4 days under WL and then transferred 4 more days to compartment plates treated with Mock, IAA ($10^{-6}$ M), tZ ($10^{-8}$ M) or both IAA and tZ, on the shoot compartment and under White Light (WL) or White Light + Far-Red light (WL+FR, n>15). Different letters depict statistical differences according to a Kruskal-Wallis test ($p<0.05$).

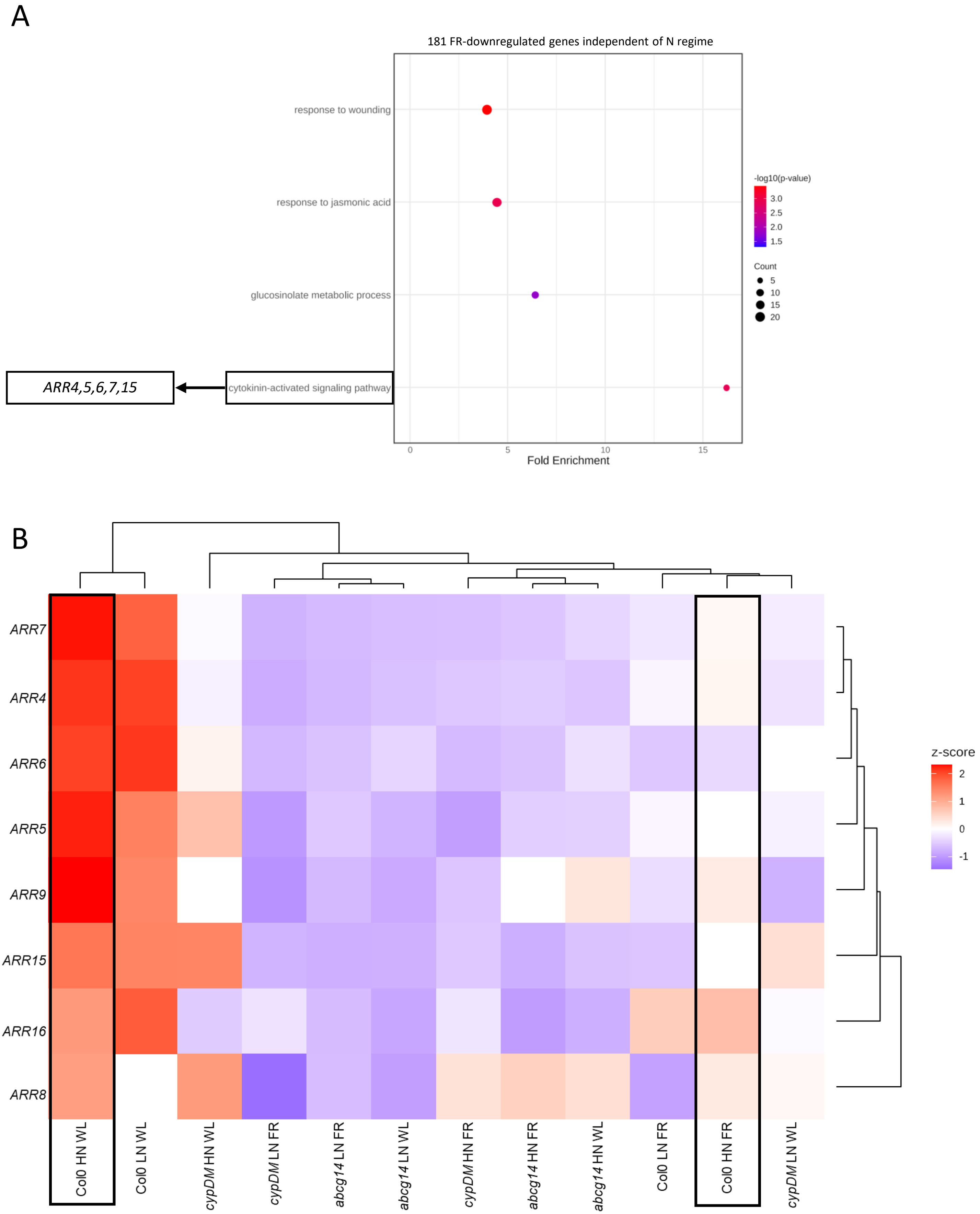


Sup. Figure 7: Expression of several type-A *ARRs* is downregulated by WL+FR

Sup. Figure 7: Expression of several type-A *ARRs* is downregulated by WL+FR

(**A**) Bubble plot representing Gene Ontology (GO) enrichment analysis for the 181 genes that are commonly downregulated by WL+FR in both Col-0 HN and LN. For GO Biological Process (BP), the most specific category subclasses with a significant enrichment (Fisher's Exact test, Bonferroni corrected, $p<0.05$) are plotted. X axis represents the fold enrichment compared to a random sample of genes. $-\log_{10}$(p-value) is indicated by colours and number of genes per category is indicated by the dots size. GO "cytokinin-activated signalling pathway" is highlighted by a black box. (**B**) Heatmap representing the z-score (indicated by colour) of detected type-A *ARRs* across all transcriptome samples. Samples and genes are clustered by similarity of regulation. The conditions "Col-0 HN WL" and "Col-0 HN FR" are highlighted with a black box. *ARR4, ARR5, ARR6, ARR7* and *ARR15* are significantly different between those two conditions and show the largest dissimilarities ($\log_2$ FC≥1, FDR<0.05).

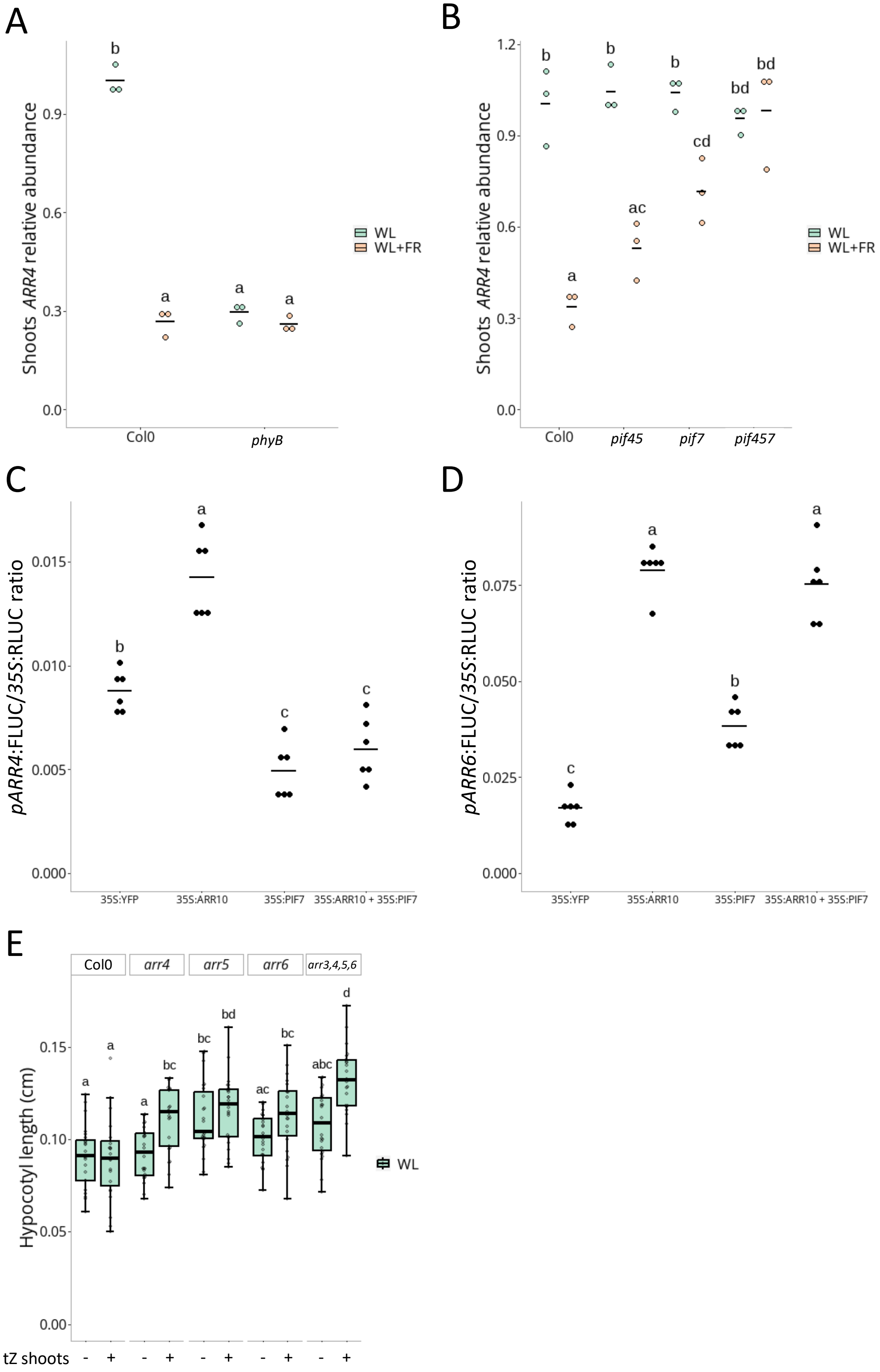


Sup. Figure 8: WL+FR elicits transcriptional downregulation of type-A *ARRs*, in a phyB and PIF7-dependent manner, and type-A ARRs prevent tZ-induced elongation in WL conditions

Sup. Figure 8: WL+FR elicits transcriptional downregulation of type-A *ARRs*, in a phyB and PIF7-dependent manner, and type-A *ARR*s prevent tZ-induced elongation in WL conditions

(**A**) Associated to Figure 4B. Shoots *ARR4* transcripts relative abundance measured by qPCR in Col-0 and *phyB;* or *pif45*, *pif7* and *pif457* (**B**, associated to Figure 4E). Each dot represents a biological replicate (pool of n>20 plants) and black bars the mean of the biological replicates. (**C**) Transactivation assay in *Nicotiana benthamiana*. A construct expressing *pARR4:FireflyLUC;* or *pARR6:FireflyLUC* (**D**), and *p35S:RenillaLUC* was co-infiltrated with a construct expressing *p35S:YFP* (baseline control), *p35S*:*ARR10, p35S:PIF7* or both *p35S:ARR10* and *p35S:PIF7*. The FireflyLUC reporter activity was expressed ratiometrically to the RenillaLUC internal control. Each dot represents a biological replicate (n=6) and black bars the mean of the biological replicates. Different letters depict significant differences according to a two-way ANOVA followed by a Tukey's post hoc test ($p < 0.05$). For qPCR and transactivation experiments (**A**-**D**), different letters depict significant differences according to a two-way ANOVA followed by a Tukey's post hoc test ($p<0.05$). (**E**) Hypocotyl length in cm of Col-0, *arr4, arr5, arr6* and *arr3,4,5,6* seedlings grown for 4 days under WL and then transferred 4 more days to compartment plates treated with Mock or tZ ($10^{-8}$ M) on the shoot compartment (n>21). Different letters depict statistical differences according to a Kruskal-Wallis test ($p<0.05$).
